# TiFM2.0 – Versatile mechanical measurement and actuation in live embryos

**DOI:** 10.1101/2024.11.10.622720

**Authors:** Ana R. Hernandez-Rodriguez, Yisha Lan, Fengtong Ji, Susannah B.P. McLaren, Joana M. N. Vidigueira, Ruoheng Li, Yixin Dai, Emily Holmes, Lauren D. Moon, Lakshmi Balasubramaniam, Fengzhu Xiong

**Affiliations:** Gurdon Institute, University of Cambridge, Cambridge, United Kingdom; Department of Physiology, Development and Neuroscience, University of Cambridge, Cambridge, United Kingdom; Life Science Institute, Zhejiang University, Hangzhou, China

## Abstract

During development, spatial-temporally patterned tissue-level stresses and mechanical properties create diverse tissue shapes. To understand the mechanics of small-scale embryonic tissues, precisely controlled sensors and actuators are needed. Previously, we reported a control-based approach named tissue force microscopy (TiFM1.0), which combines dynamic positioning and imaging of an inserted cantilever probe to directly measure and impose forces in early avian embryos. Here we present an upgraded system (TiFM2.0) that utilises interferometer positioning to minimise probe holder footprint, enhancing accessibility and imaging signal. This new design enables a double-probe configuration for bidirectional stretching, compression and stress propagation experiments. As proof-of-concept, we showcase a variety of examples of TiFM2.0 applications in chicken and zebrafish embryos, including the characterization of mechanical heterogeneities important for the morphogenesis of the chicken posterior body axis. We also present simplified designs and protocols for the replication of TiFM systems with minimal custom engineering for developmental biology labs.

## INTRODUCTION

Multicellular organisms create tissue shapes throughout development by actively regulating mechanical forces and tissue material properties through patterned cell activities^1–6^. These mechanical factors also provide feedback to cell behaviours and genetic regulation, forming a close interplay that contributes to the robustness of morphogenesis^7–11^. Such interplays are deeply embedded in the evolution history of development and work across scales, leading to a high degree of complexity. Disentangling these complexities would require quantitative measurement and control of tissue mechanics, integrated with molecular and cellular approaches, such as -omics^12^ and high-resolution live imaging^13^. These precision mechanical tools are also expected to impact engineering efforts that aim to recreate tissue and organs of functional shapes^14^.

Existing *in vivo* tissue mechanics approaches can be categorised largely by the presence or absence of direct contact with the sample. Non-contact methods include light and acoustic imaging of the sample and interpretation of the returned signal under certain assumptions, such as tissue geometry, composition and viscoelastic behaviours. The signals range from the expression and dynamics of key cytoskeleton-related molecules^15,16^, their associated sensors^17^, cell shapes and movements^18,19^, cell/tissue recoils triggered by laser ablation^20^ or optogenetics^21,22^, to scattering patterns and spectrum shifts^23–25^. These approaches offer accessibility with low-invasiveness and can cover large tissue areas. The disadvantages are that they often do not yield quantitatively definitive results as the measurements usually capture only a fraction of the factors that contribute to the tissue forces and properties, and assumptions used in data interpretation can strongly bias the conclusions. For example, inferring junctional tension and pressure from image analysis often assumes equilibrium conditions and does not consider the 3D nature of biological tissues^26^. On the other hand, contact methods utilise an inserted or injected sensor, usually with known and/or controllable mechanical properties, to directly detect its interactions with cells and tissues. The signals recorded can be directly imaged deformations and deflections^27–31^, sometimes also measured through other means such as electrical currents^32^. Whilst providing definitive measurements enabling the establishment of the regimes of forces and mechanical properties in which developing tissues operate, the results from these approaches must be interpreted carefully, considering the effective spatial-temporal resolution and scales. In particular, whether the scales are relevant to tissue morphogenesis and how the invasiveness of sensors to the local tissue environment causes mechanical artefacts. Furthermore, considerable probe design and precision engineering expertise^27,29,33–35^ is often required in developing and deploying mechanical sensors for small and soft embryonic tissues, reducing the throughput and applicability of such systems for wide use in the developmental biology community.

The prototype TiFM system (TiFM1.0)^29^ is a cantilever-based measurement and actuation tool that vertically inserts a soft atomic force microscope (AFM) probe into intact, live embryonic tissues. The probe deflects under the morphogenetic forces from the tissue or imposed forces by the motorized holder. Real-time, precise measurements of the positions of the probe tip and holder allow the force to be quantified and controlled dynamically. TiFM1.0 used charged capacitors to monitor the position of the cantilever holder, which results in a large probe head that limits sample illumination. The capacitors are also sensitive to the humidity in the environment of the embryo sample, where condensations could cause discharges that affect the measurements. Another challenge associated with limited access space on top of the sample is that further modifications, such as multiplexing of the probes, are not allowed. Here we overcome these challenges by incorporating interferometer-based positioning, greatly reducing the probe head footprint. This new design allows for improved top illumination, a stage mounted incubation chamber, and a double-arm probe holder to be added. These modifications expand the applications possible to include different types of stress loading and rheological assessment, and enhanced coverage and reliability to provide a new generation of tissue force microscopy (TiFM2.0), suitable for studying a variety of tissue mechanics questions in live embryos.

## RESULTS

### Design and operation of the TiFM2.0 system

TiFM2.0 (Figure 1A) offers a modular design, utilizing a probe head containing either 1 or 2 electrical piezos in parallel, enabling independent and fine positioning of the probes along the X axis through a voltage controller (Figure 1Bi,ii). The whole probe head is adjustable by a 3-axis micromanipulator, while the second piezo has an additional 2-axis (YZ) micromanipulator for fine alignment of 2 probes relative to each other. The overall much smaller probe head construct (compared to TiFM1.0^29^) can be lowered into an environmental well fitted with a controlled heating holder and a water supply micropipette, where standard glass-bottom sample dishes fit. To maximize the flexibility and range of interactions in the dual-probe setting, we designed the holder for AFM chips on an inclined surface (Figure 1Bii). The inner surface (between two holders) has an angle of 11.3° to the vertical plane and the outer surface has a 20° inclination to the vertical plane. This allows the distance between two AFM chips to vary between a few hundred µm on the negative (*i*.*e*., the two probes crossing each other) to >1mm which provides wide operating space for different tissue sizes and functions including stretching, compressing and twisting. The mounting surface of the holder supports a mirror for the interferometer lasers to measure probe holder positions from the sides. The probes and sample are imaged from the bottom side via a wide-field fluorescent microscope with a high framerate camera (Figures 1C-D). Similar to TiFM1.0, the theoretical force sensitivity limit is at the order of 10nN, set by the resolution of probe tip tracking currently at the order of 1µm and the softest probes we tested at the order of 0.01N/m. In practice, we detect most embryonic tissue forces between 100nN and 10µN (tissues yield at the higher end). The micromanipulators can be further automated for programmable probe alignments, and the microscope can be changed to a high-resolution inverted confocal microscope or a fast oblique plane light-sheet microscope (OPM)^36,37^ depending on the applications required.

**Figure 1.**
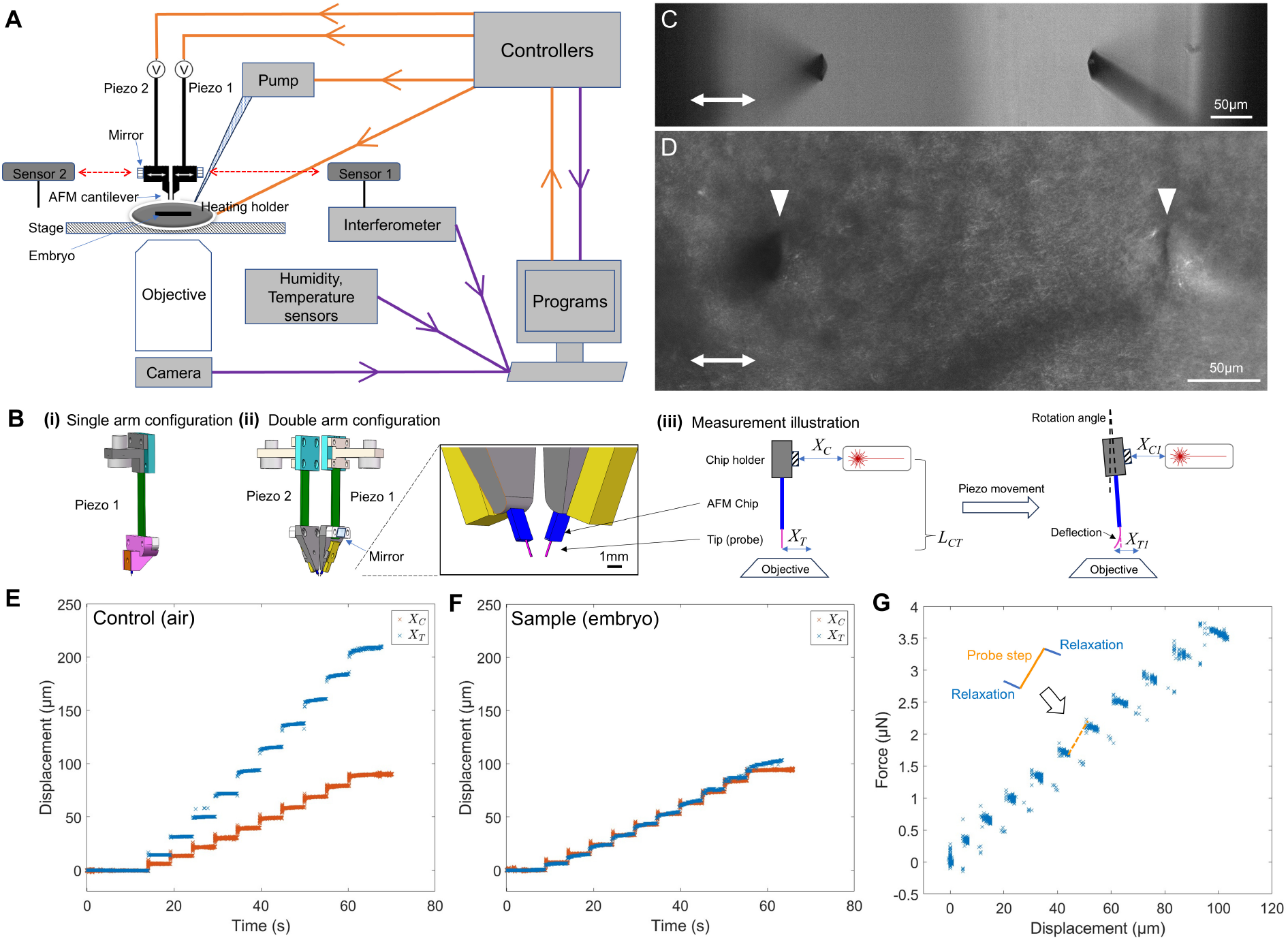
TiFM2.0 design. (A) Schematic of TiFM2.0 highlighting the design logic and main components. (B) Drawing of the single (i) and double (ii) motor probe holder. The blue probe chip and the pink cantilever probes are drawn in exaggerated sizes for illustration purpose. (iii) Illustration of measurement pipeline. *L*_CT_ marks the distance in z-axis between probe tip and the interferometer target mirror. Because of the rotational nature of the piezo, *x*_T_ will increase more than *x*_C_ under no-load as the system moves in one direction, at a ratio given by (*L*_CT_+*L*_P_)/*L*_P_, where *L*_P_ is the length of the piezo. The rotational angle of the piezo is drawn in exaggerated magnitude for illustration purpose. The z-axis movement of the probe tip resulted from this small rotation is negligible. (C) Example of double probes aligned in Z, microscope view in air (no samples), arrows indicate direction of probe movement. (D) Example of double probes inserted in an embryonic tissue (HH4 epiblast). (E) Example of a force loading control run in air. The holder was driven to make stepwise movements detected by the interferometer (*x*_*C*_), the images provide tracking of the probe tip (*x*_*T*_). (F) Example of a force loading experimental run in an embryo (~HH11, medial-to-lateral loading on the body axis), following the control in panel E. Reduced probe movements at each step compared to the trace in panel E indicate sample resistance. (G) Force estimation by comparing data in panels E and F. Stress relaxation is detected by the creep movement of the probe after each step change as a gradual reduction of measured force. See Figures S1A-F for additional examples.

An experiment on TiFM2 follows a set-up sequence where probes are aligned with samples in 3D coordinates and a control run is performed where probe movement (*x*_T_) together with the holder (*x*_C_) without resistance is recorded by the microscope and the interferometer (Figure 1Biii, E). Because of the rotational movement of the piezo and the difference of heights (*L*_CT_, Figure 1Biii) at which *x*_T_ and *x*_C_ are measured depending on the specific probe holder used, *x*_T_ and *x*_C_ change at different rates as a function of *L*_CT_ (Figures 1E, S1A,D). Next, a measurement sequence is performed where samples are inserted and actuated with the probe(s) under the control of a user-defined program, producing position data tracks of the holder (*x*_C1_) and the probe inside the sample (*x*_T1_) (Figure 1Biii, F, S1B,E). The normal force on the probe along the x axis at any time point is then given a by *F = k*cos*(θ)[f(x*_C1_*)-x*_T1_] (Figure 1G, S1C,F), and can be converted to stress as *σ = Fw*^*-1*^*(z-z*_0_)^*-1*^, where *k* is the spring constant, *w* is the width of the probes, *z* is the insertion depth, *z*_0_ is the upper tissue surface, and *θ* is the insertion angle relative to the z axis. Note that under the embedded configuration of the probe-tissue interface, the detected force/stress on the probe has multiple sources of origin (such as compressive deformation along the x axis and shear deformation along the y axis near the probe edge). Therefore, while the results are indicative of the forces and material properties of the tissue location, appropriate models (e.g., finite element models^38,39^) are needed to quantitatively interpret them together with the strain field around the probe measured by tracking markers and features from the images, to reveal mechanical heterogeneities and anisotropy of the sample (further discussed in Discussion section).

### Local stretching and compression of small embryonic tissues

A key technical challenge in mechanically perturbing early embryonic tissue is to apply controlled, physiologically relevant forces and deformations. Taking advantage of the adhesion between the actuating tools and the connectiveness of epithelial tissues, recent works have allowed small tensions and shear forces to be applied in a minimally invasive manner in the avian embryo^39–41^. TiFM systems’ vertical probe-tissue interface provides the additional advantage of applying forces on bulk tissues including mesenchymal ones^29^. In the example of the pre-somitic mesoderm (PSM), we performed local stretching with TiFM2.0 to compare the tissue responses in the anterior versus posterior PSM (*i*.*e*. aPSM vs. pPSM), regions known to have a transition of cell state^42,43^ (mesenchymal posterior to epithelial anterior), cell density^9,44^ (low posterior to high anterior) and extracellular matrix (ECM) enrichment^45,46^ and distribution^47^. A course of stepwise stretching along the anterior-posterior axis, followed by probe holding and relaxation by probe retrieval was performed in the mediolateral centres of the aPSM and pPSM (Figure 2A, Movie S1). The tissues showed distinct responses and dynamics towards the final holding strain. In the aPSM, micro-tearing (Figure 2A’, arrows, and Figure 2A’’’) appeared on the tissue to the inside of the probe tips, indicating tension build-up around the edges of the probes during stretching. In contrast, in the pPSM no tearing appeared (Figure 2A’’, 2A’’’’). At both locations, an increase of neural tube wall curvature towards the PSM under stretching was observed, indicating close links between the tissues that allow transmission of forces from the transverse contraction of the PSM under stretching to the neural tube. These tissue behaviours are unchanged by the presence of the endoderm which the probes go through from the ventral side to reach the PSM. We measured the normal strain dynamics of the tissue between the probes effectively as the distance increase between the 2 probes over the initial distance (Figure 2B). After each step movement of the motors, the probes in the pPSM were found to stabilize positions more quickly compared to the probes in the aPSM, which showed a more gradual creep. This difference is particularly apparent at higher strains (Figure 2B, arrows). Upon probe retrieval after 5 minutes of holding, the tissues (now tracked manually by the insertion wound sites) both started to recover from the deformation, with the aPSM shortening faster and to a larger extent over time (Figure 2C). To quantitatively compare the mechanical differences between aPSM and pPSM, we fitted the deformation recovery phase across multiple embryos after probe retrieval using a two-term exponential model which captured the fast (1-5 seconds) and the slow (20-50 seconds) timescales (Figure 2D). The model contains a residual strain term as a result of tissue plasticity. We found variability of max and residual strains across embryos, but they are consistently higher for the pPSM than the aPSM in paired measurements within one embryo (Figures 2E-F).

**Figure 2.**
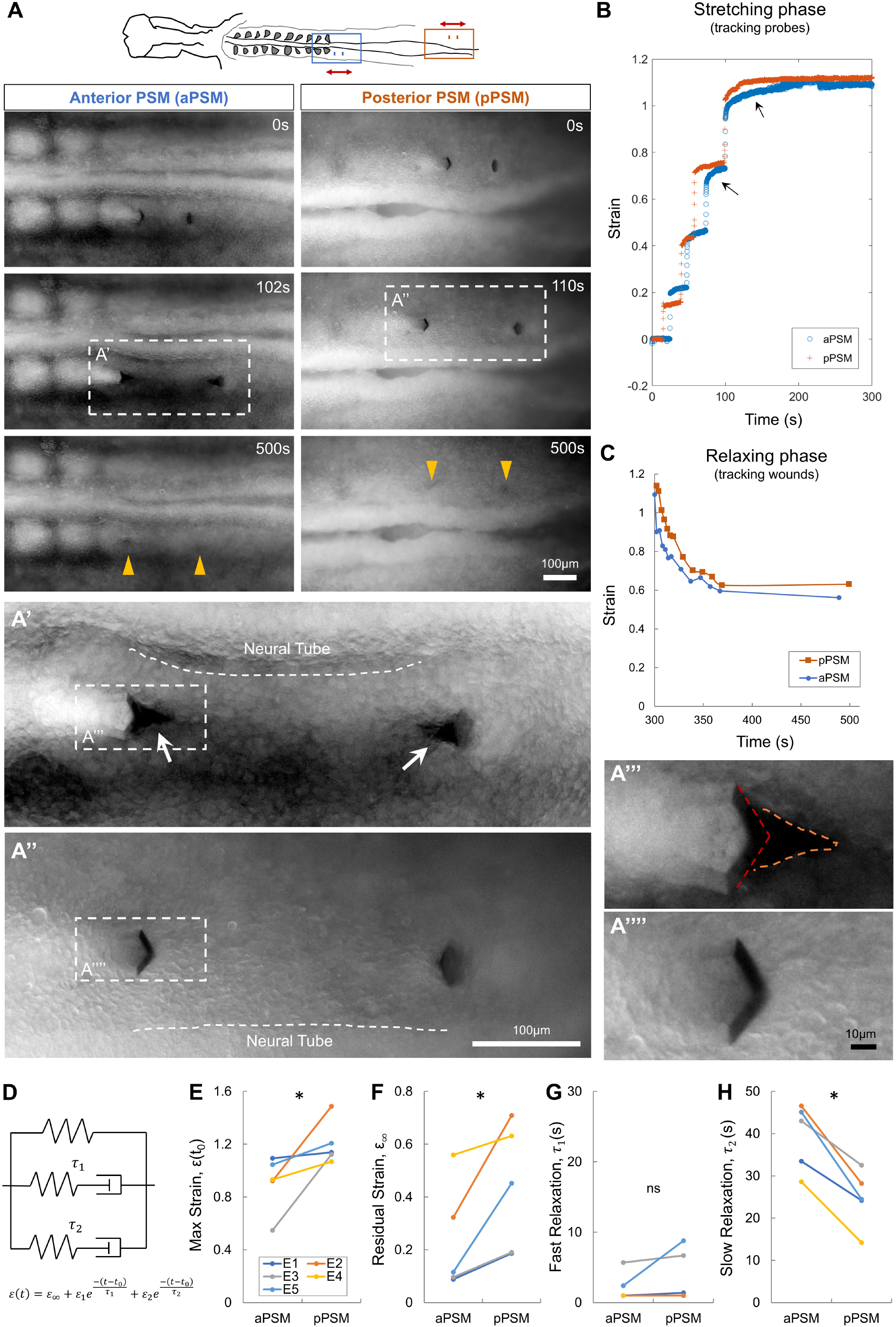
Local double probe stretching and tissue responses. (A) Stepwise stretching along the AP axis with 2 probes (*k* = 0.2N/m, Representative of N=5 different embryos, ~HH11). The schematic marks the locations in the embryo tested, arrows mark direction of probe movement. Dashed line boxes correspond to zoomed-in views in panels A’ and A’’. Yellow arrowheads track the probe wounds after probe retrieval. Dashed lines in A’ and A’’ mark the lateral side of the neural tube. Arrows in A’ indicate the tissue tears. A’’’ and A’’’’ correspond to zoomed-in views showing tissue tearing in anterior and not tearing in posterior. Dashed red lines mark the probe outline, dashed orange lines mark the tearing site. See also Movie S1. (B) Strain dynamics following stepwise stretching. Tracked from the data in panel A. The tissue length between 2 probes at each step is compared with the original length. Arrows highlight the different creep rates between aPSM and pPSM. (C) Strain dynamics following probe retraction. The timelapse movies continue from panel B. The probe wounds were tracked manually and the tissue length between 2 probes is compared with the original length. (D) Diagram of a standard linear solid model for a two-term exponential function fitted to the relaxing phase of the PSM stretching experiments. A residual strain term (ε_∞_) is included to account for the plastic deformation. (E) Initial/holding strain from experiment (comparing 5 embryos each with an aPSM and pPSM measurement, same below). Paired t-test, *p=0.029. (F) Residual strain (*ε*_∞_) from the model fit. Paired t-test, *p=0.022. (G) Fast time scale (***τ***_1_) from the model fit. Paired t-test, n.s. p=0.28. (H) Slow time scale (***τ***_2_) from the model fit. Paired t-test, *p=0.001.

The slow relaxation timescale is smaller for the pPSM, suggesting a more compliant tissue (Figures 2G-H). These simple fittings provide useful quantifications on the differences between tissue locations, but also show that simple viscoelastic models are insufficient to capture all aspects of the tissue behaviour observed. Overall, these results are consistent with the expectation from recent *in vitro* measurement^47^ that the aPSM is stiffer and more viscous than the pPSM in their native locations *in vivo*. Additionally, the deformation of tissues under the imposed forces can be analysed in terms of cell shape and arrangement changes (Figures 2A’-A’’’’). Cells can be observed to elongate along the expected tension patterns (e.g., extensive between the probes and compressive on the outer sides of the probes).

To further illustrate the potential of combining TiFM2.0 with cellular resolution live imaging, we used the epiblast of HH4 Tg(CAG:memGFP)^48^ embryos which allow single cells to be distinguished under the wide-field fluorescent microscope at 200x magnification (Figure 3A, Movie S2). We performed gradual bi-directional stretching and holding of the epiblast with live imaging, followed by segmentation of the apical surface of epiblast cells using the membrane signal with Cellpose^49^ and manual correction. Cells located in the central region between the probes show more pronounced size increase and shape changes than those of cells in the nearby off-centre region (Figures 3A-D), consistent with the region between two probes experiencing maximum tension. The shape changes show an average elongation along the direction of imposed tension (Figure 3C). To examine the dynamics of single cells during stretching, we tracked ten individual cells over time, which revealed that cell shape parameters progressively changed with variations upon stretching. These include an average apical size increase (Figure 3E), re-orientation of the long axis (Figure 3F), and alignment towards the direction of imposed tension (Figure 3G). These results show that TiFM2.0 enables cellular dynamics analysis under mechanical perturbations of the tissue.

**Figure 3.**
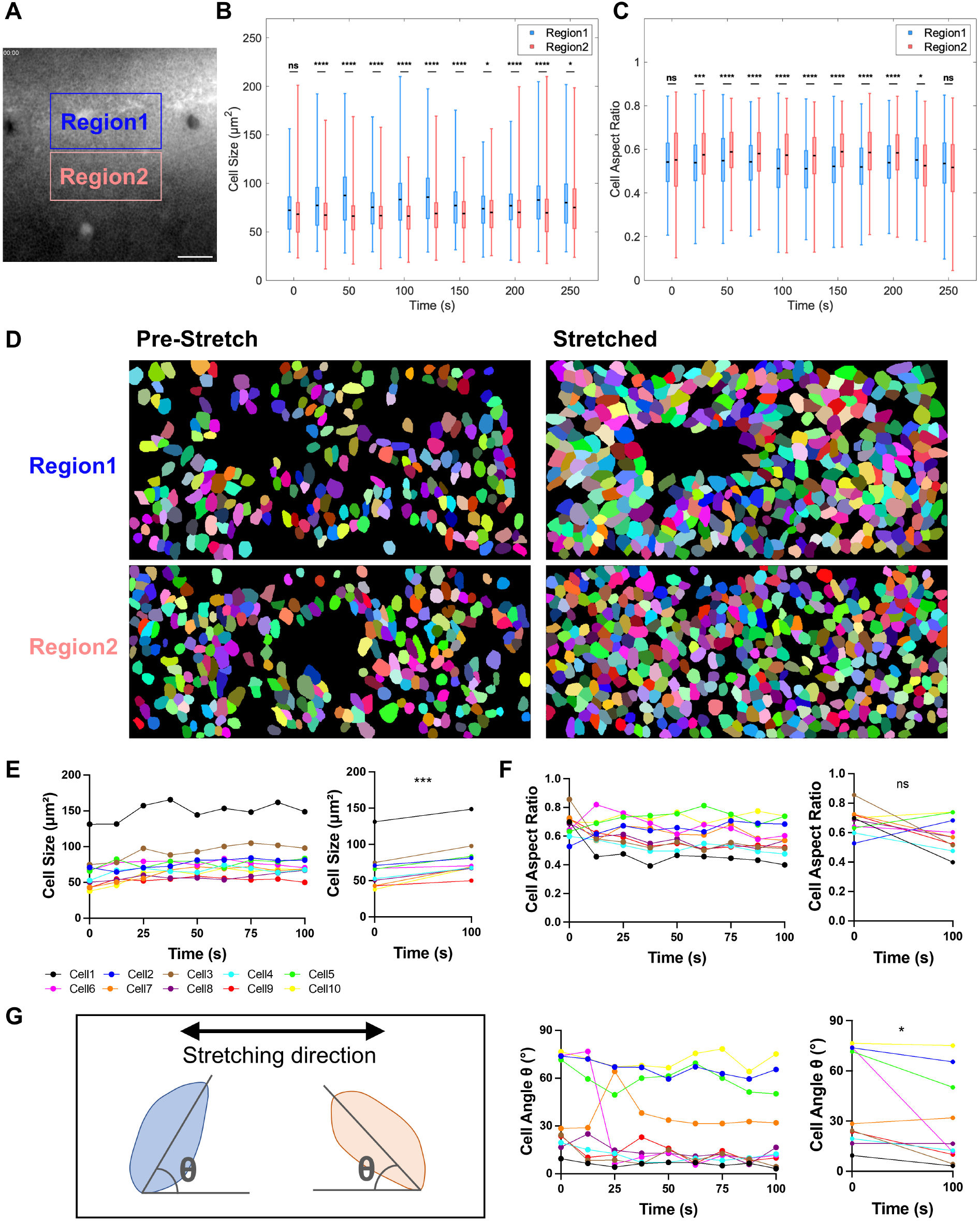
Cell geometrical changes in response to double probe stretching. (A) HH4 mem-GFP+ anterior neural plate area was imaged through bidirectional stretching (scale bar: 50µm). Two regions were selected for further analysis of cell shape changes. Region 1: located on the central line between the two probes. Region 2: positioned away from central line next to Region 1. See also Movie S2. (B) Box plot of cell size changes in the two regions indicated in (A) over time. T=0 indicates the timepoint before stretching. Within each box, black line marks mean, box extends from the 25^th^ to 75^th^ percentiles of data. In Region1, the numbers of cells included at each timepoint were 246, 263, 261, 400, 402, 405, 403, 410, 349, 222, 286. In Region2, the numbers of cells included at each timepoint were 273, 381, 409, 451, 456, 428, 387, 390, 363, 350, 309. Two-tailed Welch’s t-test at each timepoint. n.s. p=0.0693, *p= 0.0138 (t=175s), *p=0.0354 (t=250s), ****p<0.0001. (C) Box plot of cell aspect ratio changes in the two regions indicated in (A) over time. T=0 indicates the timepoint before stretching. Within each box, black line marks mean, box extends from the 25^th^ to 75^th^ percentiles of data. The numbers of cells included at each timepoint in the two regions were same as indicated in (B). Two-tailed Welch’s t-test at each timepoint. n.s. p=0.4239 (t=0s), n.s. p=0.1123 (t=250s), *p=0.0187, ***p= 0.0003, ****p<0.0001. (D) Representative example of cell segmentation masks in the two regions indicated in (A) before and after stretching. (E) Cell size changes after stretching for ten selected individual cells in Region 1. Left: cell size changes over time after stretching; Right: statistical analysis of cell size changes before and after stretching. Two-tailed paired t test (n=10), ***p=0.0001. (F) Cell aspect ratio changes after stretching for the selected cells indicated in (E). Left: cell aspect ratio changes over time after stretching; Right: statistical analysis of cell aspect ratio changes before and after stretching. Two-tailed paired t test (n=10), n.s. p=0.0932. (G) Cell angle changes after stretching for the selected cells indicated in (E). Left: Cell angle (*θ*) was defined as the angle between the cell major axis and stretching direction; Middle: cell angle changes over time after stretching; Right: statistical analysis of cell angle changes before and after stretching. Two-tailed paired t test (n=10), *p=0.0481.

To test the capacity of TiFM2.0 to modulate key morphogenetic events, we sought to promote or undo the folding process of the epithelial neural tube (Figure 4A). During neurulation, the neural plate bends dorsally towards closure at the midline to form the tube, driven by a combination of tissue-intrinsic and -extrinsic forces^1^, such as apical constriction^50^, apoptosis^51^, ectoderm tension^40,52^ and PSM compression^9^. The probes are first calibrated through synchronised control in both compressive and expansive directions (Figures 4B-C). After anchoring the probes on both sides of the neural fold edge, a sequence was performed to push the neural folds closer together and then to split them (Figure 4D, Movie S3), using the same conditions as the control sequence (Figure 4C). Comparing the probe positions in the tissue with the expected no-resistance positions from the calibration curve yields the forces at different folding states (Figure 4E). The measured forces on the probes are at the 1-10µN scale between overly-apposed to widely open neural folds, providing information of the resistance to deformation of the neural plate tissue in its native environment in the folding and unfolding directions (Figure 4F). These forces are 1-2 order of magnitude higher than recently measured native forces that drive folding, such as the compressive force from the pPSM (~100nN) measured by TiFM1.0^29^ and the net compressive forces inside the anterior neural folds (~100nN) measured by an iMeSH cylinder^53^, which drive comparatively much slower tissue deformation (order of 1hr as compared to ~10s here) during normal morphogenesis of the neural tube.

**Figure 4.**
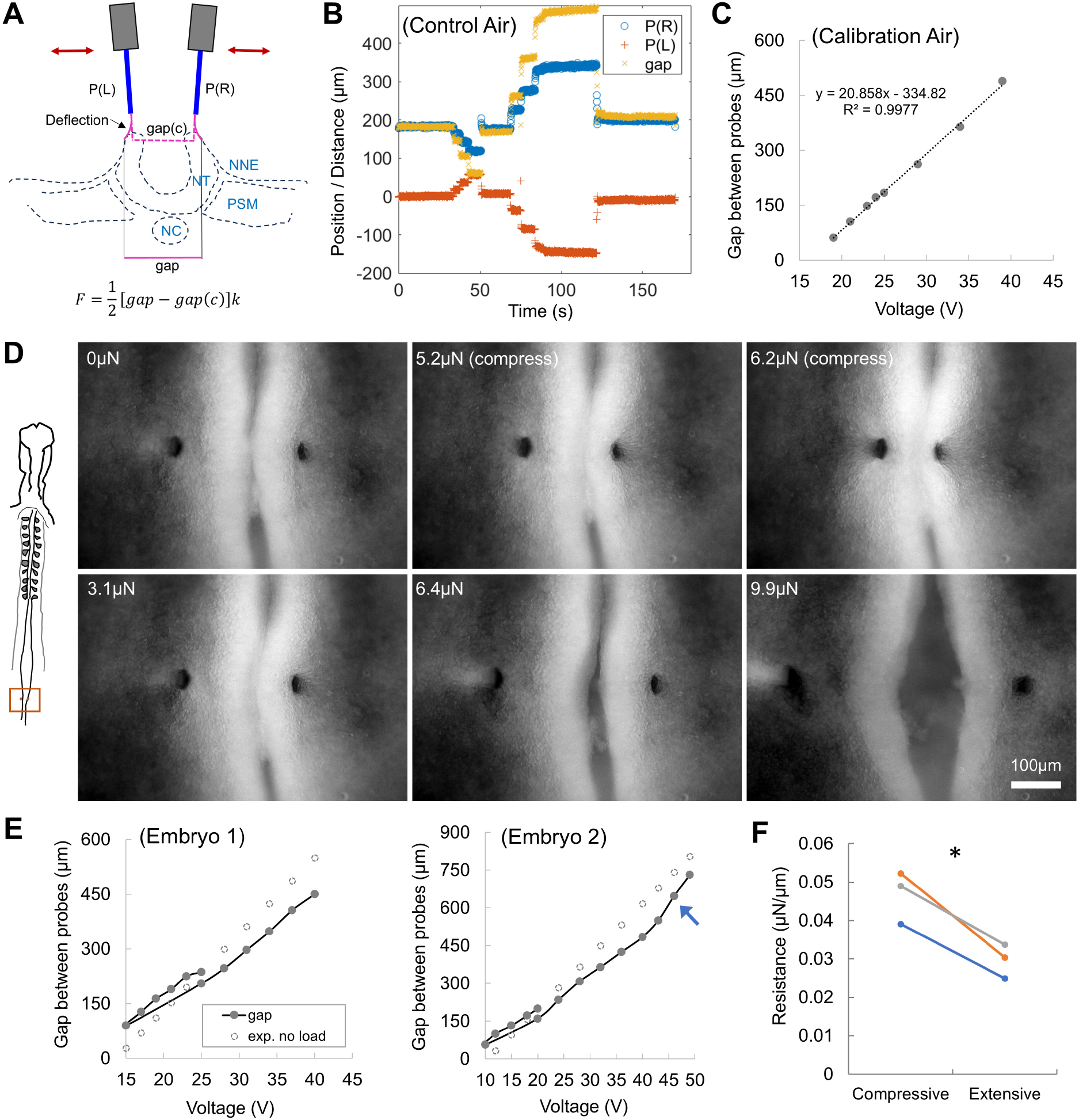
Compression and pulling of the neural folds. (A) Diagram representing experimental design. P(L) and P(R) are the synchronized left and right probes. Gap(c) is the gap between two probes in control/no-resistance condition, compared to the experimental Gap for force measurement. NT, neural tube; NNE, non-neural ectoderm; NC, notochord; PSM, presomitic mesoderm. (B) Double probe tracks from a control timelapse movie. (C) Calibration of probe gaps with the driving program to establish the no-force gap curve. (D) Images of the compressed and pulled neural folds under the driving program. Representative sample from N=4 embryos, ~HH11. The parallel bright tissues in the middle are the two neural folds. Forces labelled (in µN) represent force exerted on the probes (*k* = 0.2 N/m) by the neural folds during compression and pulling. See also Movie S3. (E) Probe gaps over the course of the perturbation for two representative embryos. Black line links the steps, starting from 25V. A roughly linear resistance increase is seen in both folding and unfolding directions. Arrow marks tissue yielding when the force exceeds the elastic regime. (F) Tissue resistance fitted from the force-displacement relationship. The neural tube shows more per unit displacement resistance in the folding direction as compared to the unfolding direction. Paired t-test, *p=0.019.

Using TiFM2.0 we can analyse the heterogeneity of tissue mechanical properties spatially, for example by pulling or compressing the posterior body axis and following its subsequent relaxation^11^. When driving the probe from posterior to anterior (A-P), we observed higher levels of tissue compression occurring near the probe (close), whilst tissues further anterior (far) show a decreased amount of deformation (Figures 5A, E). This is in contrast to driving the probe laterally from left to right (Lateral), where both close (the probe side) and far (the contralateral side) tissue regions show similar and smaller compression (Figures 5B, E). Upon probe retrieval after a 5-min holding period, the posteriorly compressed tissues close to the probe partly recover but retain a significant amount of deformation in contrast to the tissue far from the probe, which recovers a larger fraction (Figures 5C-F). In comparison, in the lateral compression, both close and far tissue regions show recovery (Figures 5C-F).

**Figure 5.**
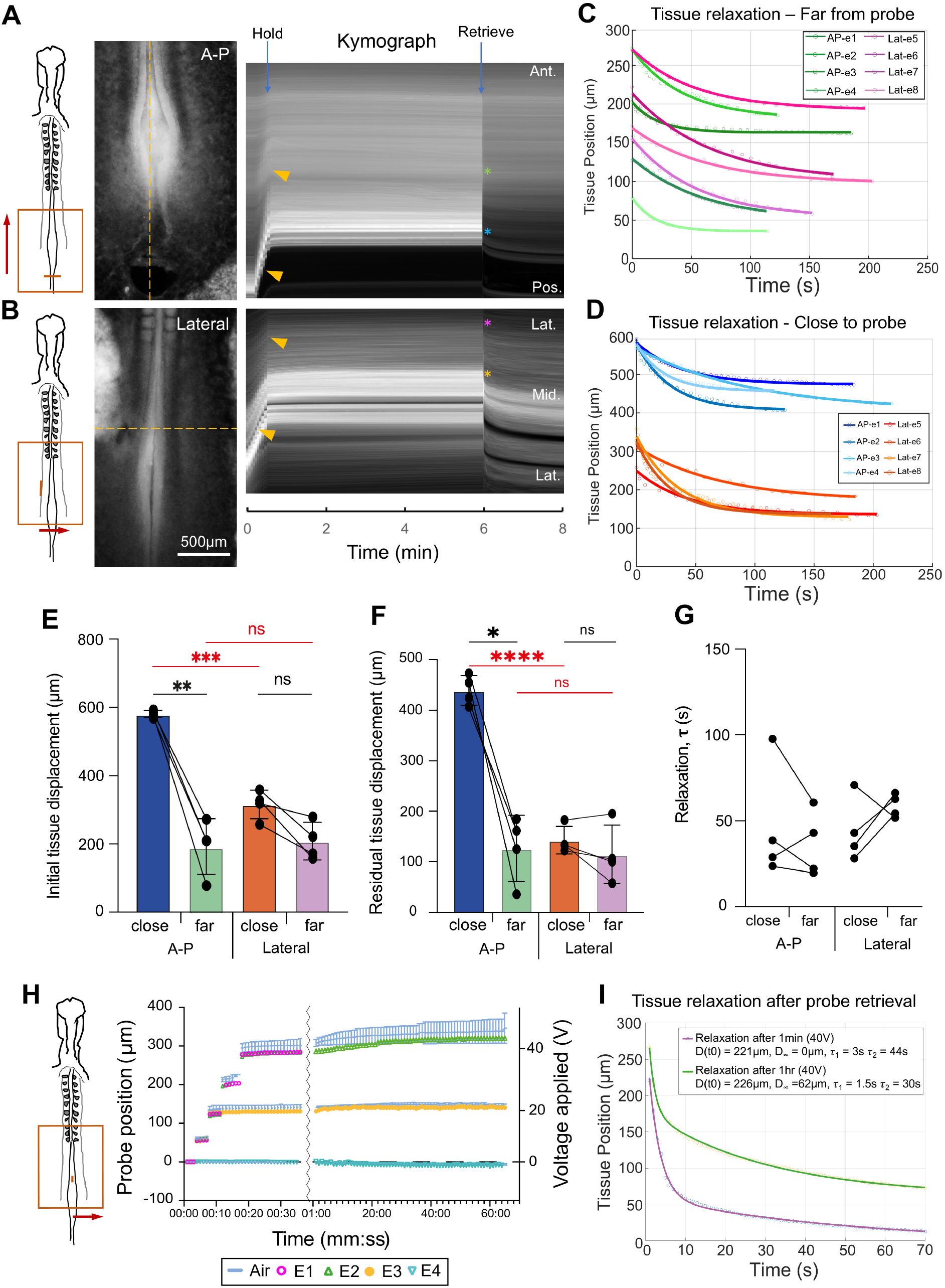
Mechanical anisotropy of AP and ML axes. (A, B) Posterior body axis anteroposterior (A) and mediolateral (B) compression using an aluminium foil probe (~400 µm wide) on ~HH10 embryos. The probe was inserted just posterior to the neural tube and moved in the anterior direction ~600 µm (A) or inserted at the PSM-LPM border and moved ~400 µm towards the midline (B). The yellow lines indicate the axes used for kymograph plotting (shown on the right). The kymograph is labelled with the points at which the maximum compression was achieved (Hold) and when the probe was retrieved (Retrieve), ~5 min 30 sec later. Arrowheads in each image compare the extent of deformation under the probe at different distances from the probe where (A) shows more drop of slope between bottom and top arrowheads compared to (B). Asterisks compare the extent of recoil close and far from the probe. (C, D) Position of tissues tracked following probe retrieval after holding compression in A-P and Lateral (Lat) directions. One term exponential decay with a residual term (line) was fitted to the deformation tracks (dots). Tracks of tissue far (C) and close (D) to probe are compared. (E, F, G) Maximum tissue displacement prior to retrieval, fitted residual tissue displacement and relaxation timescale (***τ***) after compression and hold for ~5 min in AP and Lateral compressions, close and far from the probe. Mean±s.d. In black, two-tailed paired ANOVA (E) ** p=0.0069 (F) * p= 0.0137. In red, two-tailed Welch’s t-test (E) *** p=0.0007 (F) **** p ≤ 0.0001 (n=4). (H) Mediolateral compression using notochord anchored probe (~50 µm wide) in ~HH11 embryos. Air controls show normal probe movement (n=20, mean±s.d). E1 shows compression for ~1 minute. E2 shows 40V of compression for 1 hour. E3 shows 20V compression for 1hr. E4 shows no compression for 1hr. Embryo resists probe movement. During compression, images taken at 1 second interval. During holding, images taken at 1min interval. Also see Movie S4. (I) Tissue relaxation of E1 (after 1 minute of compression) vs E2 (After 1hr of compression). Curves obtained from fitting a two-term exponential function with a residual displacement to the tracks. *D*(t0) maximum tissue displacement, *D*_∞_ residual tissue displacement, fast (***τ***1) and slow (***τ***_2_) tissue relaxation estimation. The difference in *D*_∞_ indicates plastic deformation in longer-term.

Despite the difference of the extent of recovery, the timescales are similar (Figure 5G). These results show that the developing body axis is deformable along the anterior-posterior axis while resistant along the medial-lateral axis. This pattern favours tissue elongation in a bilaterally symmetric manner. To test this mechanical pattern further in a developmentally relevant timescale, we performed the lateral bending on a cultured embryo using a notochord-anchored probe and held the deformation for 1 hour (Figure 5H, Movie S4). Unlike the 1-minute control that elastically restored most of the initial tissue shape, the 1-hour embryos exhibited a slow creep (several µm over the hour) during holding and showed significant residual deformations (order of ~50µm) after probe retraction (Figure 5H, I). This is far less than the extra elongation along the anterior-posterior axis achievable by a 1-hour pulling (~150µm) under a similar force in our previous study^29^. These findings suggest that the body axis is less deformable along the medial-lateral direction at longer timescales as well.

### Dynamic mechanical measurements

In addition to imposing forces and deformations, TiFM2.0 also enables a variety of dynamic mechanical measurements. For tissue forces, in addition to previously reported single-direction stalling stress measurement^29^, the 2^nd^ parallel probe allows pinning and clamping of tissues that are not normally held down by other structures. In the example of the early blood vessels, which lay elevated on the ventral side of the embryo above the yolk sac, the probes can be used to clamp the vessel in place and measure blood pressure (Figures 6A-D). The heartbeats are unaffected by the presence or engagement of the probes (Figure 6E). The rhythmic change of blood pressure is absorbed by the expansion of the vessel and is not detectable while the probes are not clamping onto the vessel (Figure 6C,F). During clamping, the probe tip can be observed to move with the vessel wall. The peak-valley difference varies between 5-10µm, which provides an estimate of pressure differential at 23±4Pa (n=3, probe force constant *k*=0.03N/m, and the contact area *A*=10^−8^m^2^).

**Figure 6.**
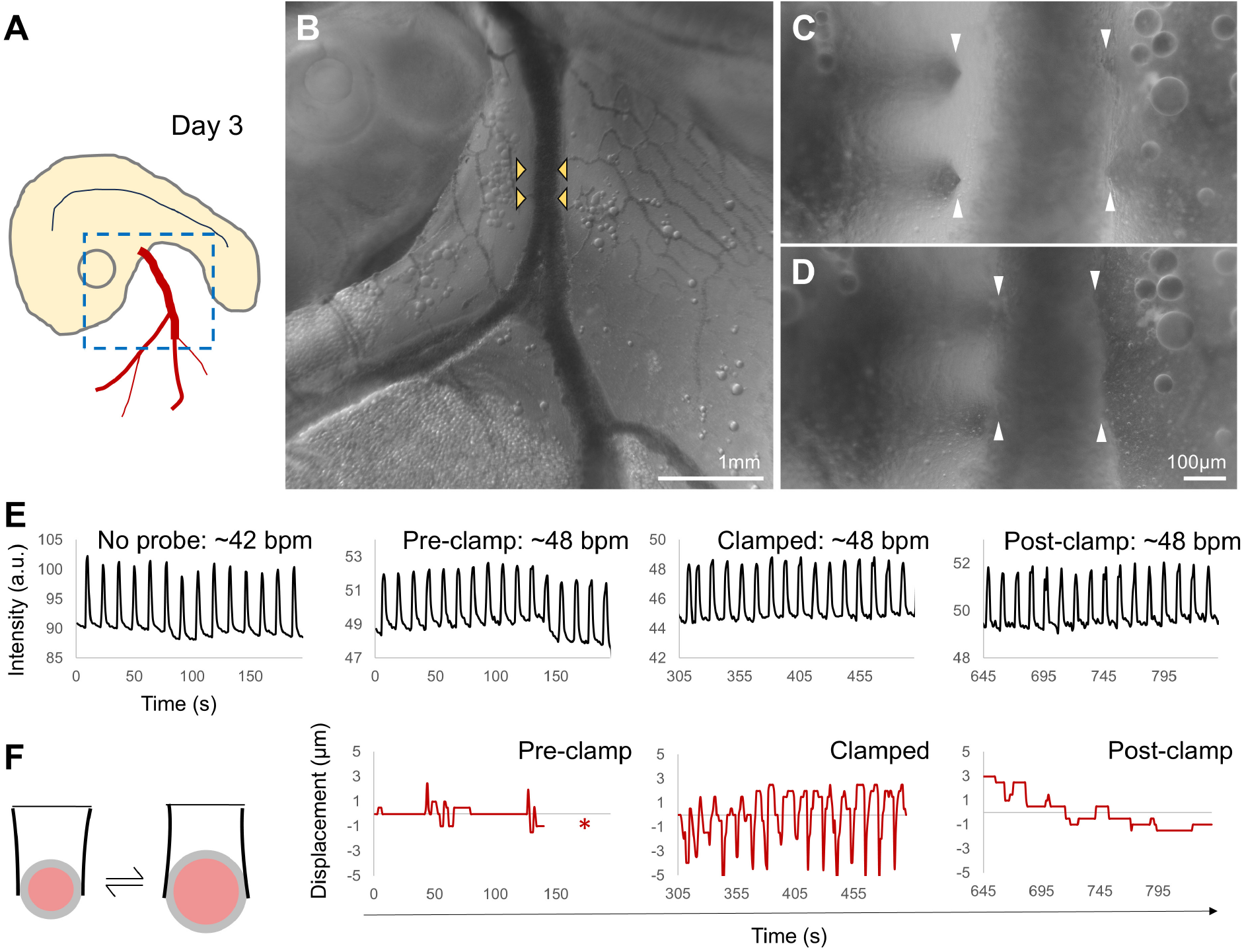
Blood pressure in the embryonic artery. (A) Diagram of a Day-3 embryo showing the imaged region and the right omphalomesenteric artery leading off the body. (B) Zoomed out image of the region and the artery. Yellow arrowheads mark the insertion points of the double-pronged probes on both sides of the artery. (C) Probes pre-clamp. The right-side probes are in contact with the artery wall. (D) Probes clamping the artery after the left-side probes are moved to engage. (E) Monitored heart rate of the embryo through the experiment. (F) Diagram and measurement of probe movement. Heartbeats send a surge of blood flow through the artery and expand the vessel, generating a transient pressure on the probes. Asterisk at the Pre-clamp plot marks the time period when the probes are being moved with no displacement measurements.

TiFM2.0 also enables *in situ* probing of tissue mechanical properties, taking advantage of accurate dynamic positioning and sensitive soft probes. The concept is that a small local motion of the probe tip inside the tissue is locally resisted and also its propagation is gradually dissipated from the source (Figure 7A). The dynamics of these deformation signals encode the viscoelastic properties of the tissue location. To test this capacity, we inserted the probes into water and a series of poly-isobutylene in hexadecane (PIB, a reference viscoelastic fluid [SRM 2490]) solutions of increasing storage modulus (Figures S1G-I). The sending probe was then driven to oscillate using a sine function with a given maximum voltage and frequency while the tip was tracked (Figures 7A-C). Measuring the amplitudes of *x*_T_ in both sending and receiving probes, we can resolve differences in the range of 300-3000Pa in storage modulus of the PIB (Figure 7Bii), a key range that spans the expected values of embryonic tissues as measured by methods introducing larger deformations^54^. At lower frequencies, we can increase the measurement sensitivity to the 100Pa range, however this sensitivity is lost at higher frequencies (Figure 7Cii). To further assess the information in the mechanical signal, we fitted *x*_T_ to a sinusoidal function and aligned the interferometer signals to the same time axis, allowing phase differences of probe movement between samples to observed. For example, the probe shows a smaller amplitude and more phase shift in 18.5% PIB compared to water (Figures S1H-I), indicating stronger viscoelastic resistance to probe movement in PIB. To test this in epithelial-tissue-like solids, we compared the amplitude of oscillation between PBS and a series of hydrogels of increasing stiffness at two different depths that are representative of experimental conditions in embryos (Figures 7D S1G, Movie S5). Overall, we observed lower amplitudes as stiffness increased in a quantitative relationship that would allow differences in amplitudes to be related back to differences in the material properties of the samples. Probes with lower force constants (e.g. *k* = 0.03N/m) can be used for samples at the softer ranges. Of note, at deeper insertions in stiffer gels, the oscillation of the probe tip is antiphase to the imposed sinewave motor motion and is represented as negative, due to the strong resistance bending the probe at a higher z-axis point. Together, these data show that TiFM2.0 can sensitively resolve the mechanical differences of viscoelastic materials in comparable ranges of the embryonic tissues. However, it is important to note that embryonic tissues in their native environment are small and anisotropic, connecting with other tissues through ECM links. The mechanical property and *x*_T_ relationships established in the homogenous control materials above cannot be directly used as calibration to estimate values at the tissue location. Extensive mapping using TiFM2.0 in conjunction with proper multi-tissue models are required to quantitatively solve tissue mechanical properties of intact embryos (further discussed in Discussion section).

**Figure 7.**
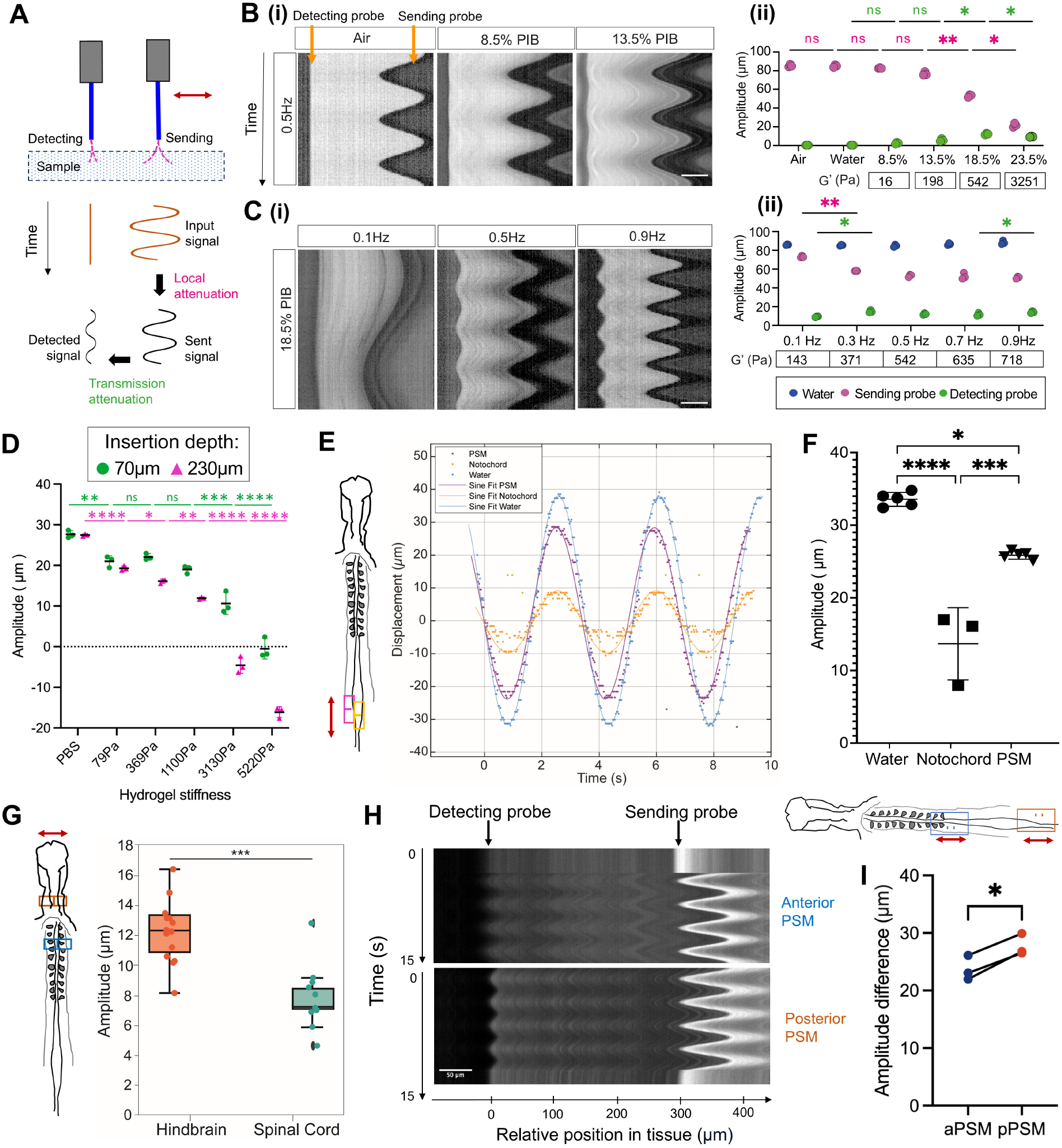
Tissue mechanical evaluation with oscillatory probe movements. (A) Diagram of oscillatory motion used to assess the mechanical properties from the displacement response. The input signal is attenuated by tissue resistance at the origin as well as through transmission. The spatial-temporal dynamics of the signal (e.g., magnitude and phase) encode viscoelastic information of the tissue location. (B,C) (i) Representative kymographs showing sending and detecting probe movement. Scale bar 50 µm. (B ii) Amplitude of sending and detecting probes (*k* = 0.5N/m) in different PIB concentrations. Input signal: 10V, 0.5Hz sinewave frequency. Probe insertion depth 70 µm (±20 µm). Tukey’s multiple comparison test (n=3). Sending probe, 13.5% vs 18.5% ** p= 0.0032, 18.5% vs 23.5% * p=0.0162. Detecting probe, 13.5% vs 18.5% * p= 0.0290, 18.5% vs 23.5% * p=0.0379. (C ii) Amplitude of water, sending and detecting probes (*k* = 0.5N/m) in 18.5% PIB at different frequencies. Input signal: 10V. Tukey’s multiple comparison test (n=3). Sending probe, 0.1Hz vs 0.3Hz ** p=0.0087. Detecting probe, 0.1Hz vs 0.3Hz * p=0.0442, 0.7Hz vs 0.9Hz * p=0.0288. G’ values of PIB samples were measured with a rheometer, see also Figure S1I. (D) Amplitude of probe (*k* = 0.2N/m) differences PBS and hydrogels of varying stiffness. Input signal: 10V, 0.5Hz. Insertion depths: circle 70 µm and triangle 230 µm. Negative amplitude is defined as tip movement in the opposite direction of the driving motor. Black line marks mean±s.d. Two-tailed Tukey’s multiple comparison test (n=3). At 70 µm: n.s 79Pa vs 369Pa p = 0.9685, n.s 369Pa vs 1100Pa p = 0.3101, ** p=0.0051, *** p=0.0008, **** p<0.0001. At 230 µm: * p=0.0258, ** p=0.0037, **** p<0.0001. See also Movie S5. (E) Representative example of probe tip position tracking (marked by dots) and sin fitting (marked by line) across Water, PSM and notochord measurements (HH9-10 embryos). Sine fitting over n=2.5 periods. See also Movie S6. (F) Amplitude differences between Water (n=5), Notochord (n=3) and PSM (n=5) in HH9-10 embryos. Probe force constant *k* = 0.2N/m,10V, 0.5Hz. Mean±s.d shown. Tissue insertion depth = 70µm. Two-tailed Tukey’s multiple comparison test: * p=0.0152, *** p=0.0007, **** p< 0.0001. (G) Amplitude of probe (*k* = 0.02N/m) movement driven at 0.2Hz and 5V in the hindbrain (n=14) and spinal cord (n=10) of HH11-12 chicken embryos. Mann-Whitney U test, *** p<0.001. (H) Kymographs representing double probe movement tracks for anterior (aPSM) and posterior (pPSM) locations. Representative of N=3 samples, HH10 embryos. Sending and detecting probe amplitude tracked by high contrast lines. *k* = 0.03N/m, 10V, 0.5Hz. Scale bar = 50µm. See also Movie S7. (I) Amplitude difference between sending and detecting probes in anterior PSM (aPSM) and posterior PSM (pPSM). A smaller amplitude difference shows the detecting probe oscillates more in response to the sending probe, suggesting the tissue is stiffer. A large amplitude difference indicates the detecting probe does not respond to the imposed oscillation from the sending probe, suggesting a softer tissue. Two-tailed Paired t-test *p=0.0176, n=3.

Nonetheless, TiFM2.0 is readily available to assess developmentally-important mechanical property patterns *in vivo* and *in situ*. Using the oscillatory input as described above, we tested distinct chicken embryo body axis tissues following the parameters of Figure 7D. The notochord shows a strong restriction on probe movement, whereas PSM is much less resistant (Figures 7E-F, Movie S6). In the dorsal neural tube, the hindbrain shows less resistance to probe movement compared with the spinal cord (Figure 7G). These results are consistent with previous studies assessing the deformability of these tissues using other methods^55,56^. However, it should be noted that keeping track of the spatial range of tissue deformation by the probe is important for the interpretation of these results. For solid tissues, the resistance to probe movement measured could indicate the neighbouring tissue connections and boundaries rather than local properties around the probe, which can be much stronger. Using the second probe helps provide a defined spatial range. We tested this in the PSM where small mechanical differences between the anterior and posterior exist, as shown by stretching experiments (Figures 2B-H). Indeed, the mechanical wave propagates with a smaller magnitude loss in the aPSM as compared to the pPSM, indicating that aPSM is more solid-like while pPSM is more liquid-like relatively (Figures 7H-I, Movie S7). These results are consistent with the reported cell and ECM organisation of the PSM along the AP axis^46,47^, and recent magnetic-droplet-based rheological measurements in the zebrafish PSM^57^, showing conservation of the PSM mechanical property gradient among vertebrates despite distinct tissue sizes and ECM organisations. Together, our results demonstrate TiFM2.0’s capacity for direct *in vivo, in situ* tissue mechanical property assessment.

To expand the capacity of TiFM2.0 beyond the thin, flat avian early embryo tissues, we extended the back end of the probe with a DiI bead (Figure 8A) which provides high-contrast fluorescent signal and additional spatial coverage when the probe front becomes obscured by thicker or less transparent tissues. Using zebrafish blastula-stage embryos, which have a spherical geometry with the yolk and consequently form a thick tissue to image from the side, we tapped the trailing probe on the animal pole tissue, using fish medium and a later somitogenesis stage embryo as controls (Figures 8B-C, Movie S8). Shortly after probe contact with the animal pole the front becomes obscured by the tissue and can no longer be tracked but the extended dye is clearly trackable (Figure 8C). This allows an accurate determination of the animal pole blastoderm’s deformation to the probe (Figures 8D-F), which suggests that it is softer than the dorsal tissue of the later stage embryo, as evidenced by the consistently larger probe displacements observed at each loaded voltage (10V, 30V and 50V). These experiments show the feasibility of TiFM2.0 for side measurements of thicker tissues and a different model organism following probe modifications.

**Figure 8.**
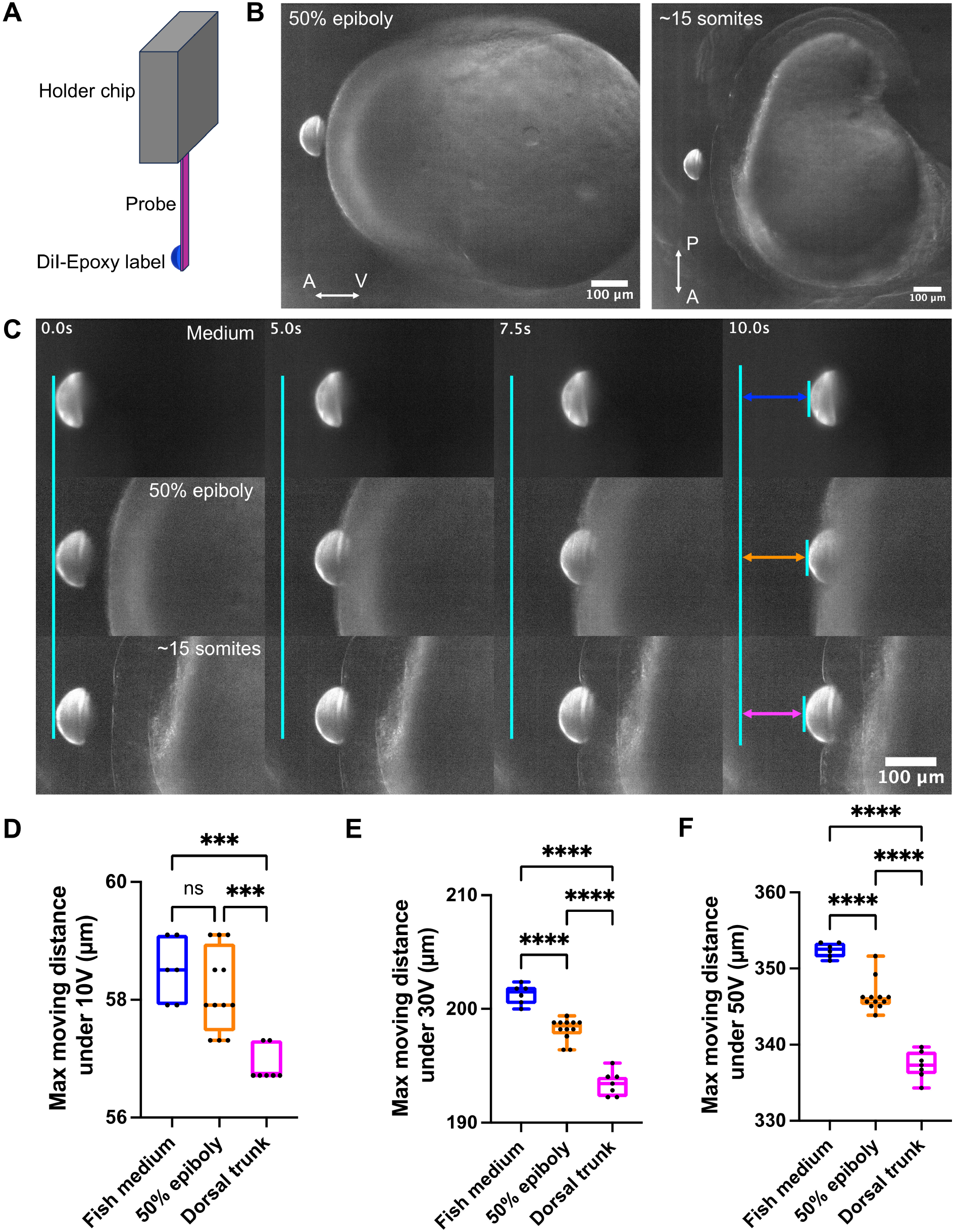
Tracking modified probe in thick tissues in zebrafish embryos. (A) Diagram of the probe with a DiI-Epoxy label. (B) Example images of the labelled probe and the sample embryo. The animal pole at 50% epiboly (left) and the dorsal trunk during somitogenesis (right). Movies are taken with red fluorescence and a low level of background light to visualize the embryo tissues. A-V axis indicates animal-vegetal; P-A axis indicates posterior-anterior. (C) Imaging of the probe movement when pushing different tissues under the voltage loaded on the probe of 20V, 0.1 Hz and Ramp function (linear movement of the motor). Fish medium (1st row), animal pole at 50% epiboly (2nd row) and dorsal trunk at ~15 somite stage (3rd row) are shown here as representative examples. Images show the probe position at the beginning of loading the voltage (1st column), starting to touch the tissue edges (2nd column), continuing to push the tissues (3rd column) and the final position when the maximum voltage is reached (4th column). The blue, orange, and magenta arrows indicate the maximum moving distance of the probe under pushing in fish medium, epiboly, and dorsal trunk, respectively. See also Movie S8. (D) Resistance comparison (shown by maximum moving distance of the probe, tracked by the bright edge kymograph of the backend of the probe, same as panel E and F) of fish medium, animal pole at 50% epiboly and dorsal trunk at 10V maximum displacements. Ordinary one-way ANOVA multiple comparison test. Fish medium (n=6) vs 50% epiboly (n=12), n.s=0.4581. Fish medium (n=6) vs dorsal trunk (n=7), ***p=0.0001. 50% epiboly (n=12) vs dorsal trunk (n=7), ***p=0.0003. (E) Resistance comparison of 30V maximum displacements in the same embryos shown in (D). Ordinary one-way ANOVA multiple comparison test, ****p<0.0001. (F) Resistance comparison of 50V maximum displacements in the same embryos shown in (D). Ordinary one-way ANOVA multiple comparison test, ****p<0.0001.

## DISCUSSION

In this work we provided proof-of-concept demonstrations of the versatility of TiFM2.0 in probing and perturbing tissue mechanics in early vertebrate embryos, with precision, flexibility, and tissue-depth coverage at biologically relevant timescales and forces. These experiments can reveal new biological insights, in combination with other mechanical tools, imaging and molecular approaches^11,56^. Notably, the collection of evidence so far starts to constitute a new level of understanding on the tissue mechanics of the avian embryonic posterior body. The tissues are produced at the posterior end and follow a posterior to anterior differentiation gradient that is now also revealed to be in parallel with a mechanical property gradient. The PSM cells, upon exiting the progenitor domain and entering the PSM, simultaneously drive tissue expansion and softening as they exhibit high motility and reorganize the ECM to absorb water^11,46,47^. This orients PSM deformation towards the posterior, and promotes the deformation of axial tissues such as the neural tube towards narrowing (including convergence and folding) and extension^9,29,58^. The stiffer axial tissues push through the “melting” progenitor domain (undergoing epithelial-mesenchymal transition^59^) containing new PSM cells, bisecting them to supply the bilateral sides^9^. These mechanical patterns and interactions enable directionality^46^, tissue coupling^9^ and a robust speed^11^, as well as potentially other key features of the body axis such as bilateral symmetry. TiFM here provides important direct measurements and controlled perturbations that put this description into proper physical context.

An important next step is to quantitatively interpret mechanical patterns and heterogeneities in the embryonic tissues uncovered by TiFM2.0. The *in situ* probing of tissue mechanics at the scale relevant for morphogenesis (~30-100µm) faces the challenge of an inhomogeneous and complex surrounding that in many cases can make the source of stress/resistance to the probes unclear, as early embryonic tissue sizes are comparable to the scale of measurement. Physically separating different tissues (such as by microsurgery) that could contribute to the probe dynamics would likely disrupt key ECM links between tissues and introduce mechanical effects at the wounds (such as tension increase), limiting the effectiveness of experimental dissection of different contributors. To address these challenges, a multi-tissue model that allows systematic fitting of TiFM data to the complex tissue environment is needed. To start, the complex modulus of the control materials will be recapitulated with the probe geometry and measurement dynamics.

This will allow characterization of the strain field around the probe in the materials. Next, the model will be extended by supplying inhomogeneous samples, such as two connected but distinct tissues (e.g., PSM and neural tube) where assumptions are made about their material properties and their ECM connections. Experimental data, such as mechanical signal propagation along and across tissues, conducted systematically at different depths, frequencies, directions and locations will then be used to fit the parameters of the model. Finally, TiFM actuation will be deployed to directly test, for example, the strength of ECM connections between tissues. These approaches may benefit from an experimental model of intermediate complexity, such as multi-tissue body axis organoids^60^ created *in vitro*. These steps will establish a comprehensive picture on how tissue properties coordinate complex morphogenesis and realize the full potential of TiFM systems. The avian embryonic body axis here provides an excellent model for this type of systematic deconstruction of mechanical interactions. The understanding gained will illuminate human development during these stages when many defects arise, and where comparable processes have not been described due to the scarcity of embryo samples. More generally, the rationale and approaches of TiFM applied here to the body axis model are widely relevant to other classes of fundamental morphogenesis processes.

While TiFM2.0 as described in the current work contains a variety of modules and a complex user protocol, the core mechanism and function of TiFM systems as a controlled cantilever are simple^29,39,41^, and can be deployed in a compact, easy-to-construct version (see methods – simple TiFM) or upgraded to work with high resolution microscopy, suited for large tissue-level measurement and perturbations or dynamic observation of cellular and molecular changes associated with tissue forces, respectively. Moreover, the system can be combined with other mechanical modules such as tension^40^ and magnetic^27^ controllers to enable multiplexed mechanical measurement and perturbation. Continued innovations in probe design and positioning as well as theoretical modelling for data quantification will further improve the applicability of TiFM systems and reduce their footprint and cost, enabling potentially wide use in developmental biology research and other settings.

## Author Contributions

F.X. designed the project and constructed the hardware. A.R.H. developed the measurement protocols and the incubation module; Y.L. developed the trailing probe label and performed all zebrafish experiments; A.R.H., Y.L. and F.X. performed most calibration, standard material and chicken embryo experiments, created software and analysed the data; F.J. designed and created the double probe holder; F.J., S.B.P.M., J.M.N.V., Y.D., R.L., E.H., L.D.M. and L.B. contributed to conceptualization, experiments, protocols and analysis tools. A.R.H., Y.L. and F.X. prepared the manuscript.

## Acknowledgement

The authors thank Chon U Chan and members of the K. Kawaguchi lab for suggestions on the system design and data analysis; members of the Xiong lab for reagents, technical assistance, and comments; Chun Yuan Hii and Clare Buckley for assistance with zebrafish embryos, which were obtained through the fish facility at the Department of Physiology, Development and Neuroscience of University of Cambridge; Charles Bradshaw (Gurdon institute) and James Steele (the Maxwell Centre of University of Cambridge) for 3D printing; Michela Geri and Gareth McKinley for sharing the OWCh software and suggestions; Chiu Fan Lee for suggestion of the mechanical propagation experiment. This study is supported by a Wellcome Trust / Royal Society Sir Henry Dale Fellowship (215439/Z/19/Z) and a UKRI-EPSRC Frontier Research Grant (EP/X023761/1, originally selected as an ERC Starting Grant) to F.X. A.R.H. also acknowledges an SBS-Gurdon Studentship. Y.L. also acknowledges a Cambridge International Scholarship. F.J. and L.B. also acknowledges a Herschel Smith Postdoctoral Fellowship. F.J. also acknowledges a Wellcome Early Career Award (302541/Z/23/Z). Y.D. also acknowledges an LSI-Gurdon exchange programme supported by China Scholarship Council. R.L. also acknowledges a Xuetang summer scholarship from Tsinghua University and a Harding Distinguished Postgraduate Scholarship. L.D.M. also acknowledges a Wellcome Trust PhD studentship (222274/Z/20/Z). The authors declare no competing interest.

## MATERIALS & METHODS

### List of hardware components

An equipment and component list is provided for the TiFM2.0 system. The table is organized by different modules. A simple/minimal TiFM setup is highlighted by selective components. It is not necessary to procure identical items. In particular, accessories such as connecting parts and scaffolds should be chosen according to the configuration of the base microscope. Basic instrument construction knowledge is needed to assemble the systems.

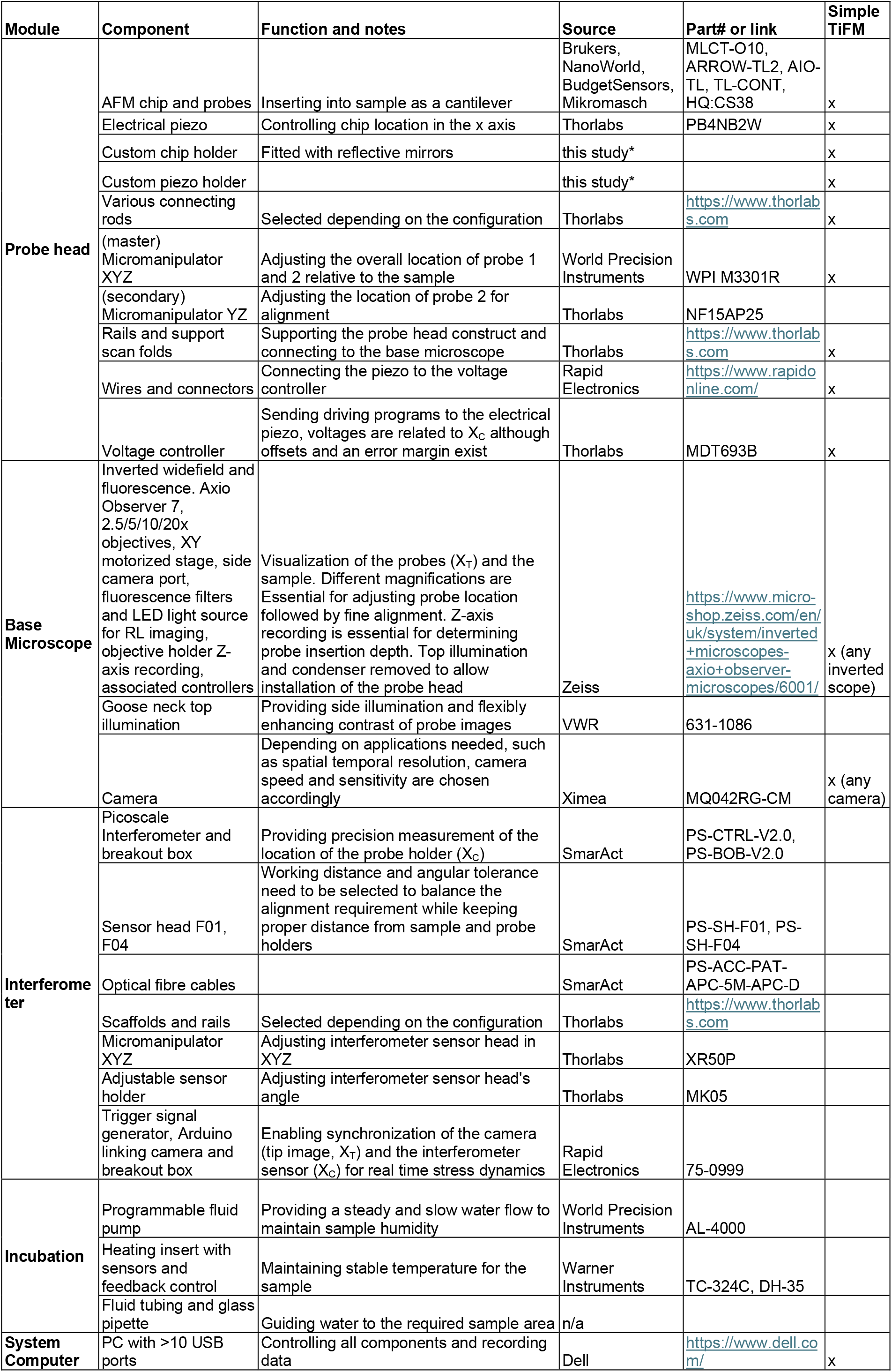

*Custom components include the motor and AFM chip holder module, and were printed by a 3D Printer (UltiMaker 3, Print Core AA 0.40) using PLA filament (2.85mm, RS PRO 832-0273). A tailored mirror (Silicon wafer, N-Phos) was glued to the outer side of the AFM chip holder for interferometer positioning. The engineering models and other details are available upon request. Custom probes were made by glueing aluminium foil (wide probe used for compression) or DiI-Epoxy beads to the end of the AFM cantilevers.

### Working environment and maintenance

TiFM systems should ideally be set up with vibration-isolation in quiet areas with stable temperature and humidity, and minimal dust. TiFM2.0’s interferometer positioning system is robust to sample humidity and does not require regular adjustment and cleaning of the capacitor sensors in TiFM1.0, but the environmental temperature and humidity should still be recorded as they can affect the sensor parameters. All wire connections should be checked regularly to ensure proper connection. The optical fibre wires should not be acutely bent or under tension. The piezo, probe holder and probes should be uninstalled from the probe head micromanipulators and stored in safe boxes when not in use as they are fragile. When installing the probe chips, a stand for the probe holder is recommended to free both hands. Under good lighting where the thin cantilever(s) is visible by naked eye, the user picks up the probe chip with a pair of sharp tweezers, holding the chip while staying away from the end with cantilevers, and insert the chip into the holder slot, before tightening with the screwdriver. When installing the probe holder to the scaffold and micromanipulators, take care to securely connect the piezo cables to the out cable from the voltage controller with the right match to avoid damaging the piezo. Upon finishing the experiments, use a clean imaging dish filled with deionized water to rinse the probe by dipping the chip/probe into the water a few times using the micromanipulator, before uninstalling the probe and holders.

### Operation protocols

When ready to use the TiFM system, power up all hardware and open all controller software on the PC. In the following Ximea Cam tool (Ximea), MT voltage controller (Thorlabs), and Picoscale interferometer software (SmarAct) are used as examples to outline the protocol, which can be adapted to other hardware and software accordingly. First, check if the acquisition systems are functioning normally by starting live imaging, and change voltage input to move the piezo. Next, prepare an embryo sample and mount it on the imaging dish as desired for the actual measurements, add a few drops of PBS/Ringers solution on top to prevent drying (thin albumen is also ok with strong probes such as 0.2N/m but may damage the weaker ones due to its adhesion and viscosity). Mount the dish on the stage, position it so that the area of interest is centered, turn on the gooseneck LEDs to provide illumination. Use 2.5x, 5x and 10x objectives to observe the embryo with the camera, adjust the lighting angles and exposure times as necessary. Next, focus on the tissue of interest under 10x, record the focus positions of the tissue surface and the depth of interest, leaving the focus unchanged for probe adjustment. Remove the sample dish and lower the scaffold holding the piezo and chip/probe, carefully approach the stage to avoid collision with the objective and centre the probe tip at the pre-determined focal plane using the XYZ micromanipulator. This ensures the tip and the tissue of interest will be aligned. Use the second YZ micromanipulator to align the second probe to the desired relative position with the first probe, usually at the same focal plane / z-axis position, if applicable.

When the probes are in position, the interferometer sensor should be aligned to the probe holders. First algin the sensor head roughly with the holder mirror using the guide laser. Then in the Picoscale software, use the “Adjustment” mode to obtain the signal pattern on screen, while adjusting the position and angle of the sensor holder to improve the alignment as evaluated by the pattern score, using the micromanipulators of the mirror mount. Leave the sensor holders untouched once a good signal is obtained (quality>60), continue with automatic adjustment and wait until the software indicates that the sensor channel is valid. Next set up the trigger signal required for the camera to activate the measurements to achieve synchronization following the software instructions. Open the data stream monitor and assign the trigger. Set up the camera to send the trigger signal upon activating frames (i.e., image acquisition). Test the trigger by starting live stream of the camera, the interferometer signal should immediately show on the monitor plot.

With probe and interferometer stand-by, the control sequence can be performed, followed by the samples. For the control run, image an empty or water sample with the moving probe under the same voltage program to be used on the embryo tissues, such as a step/ramp push for loading experiments, or an oscillatory movement for rheology experiments. Before lifting the probe holder to allow mounting of the embryo sample, mark the position of the probe tip on the screen, as it will be difficult to see the probe tip directly before it making contacting the sample, once the sample is between the probe and the microscope. This mark helps when positioning the sample to the extract desired location of probe insertion. After mounting the embryo sample, adjust the focus to determine the location of the top surface of the sample, record the Z position of the surface and the probe tip Z position after insertion. These numbers help determine the insertion depth and probe-tissue contact area. This is not required for thin tissues, where the tissue thickness and probe width determine the contact area. To insert the probe into the sample, now without a clear visual of the probe tip, lower the probe holder in Z slowly and carefully while watching the changes on the live camera stream. The probe would change the lighting seen on the live stream and cast a shadow when it enters the tissue. Adjust the focus slightly to ensure a good contrast of the probe tip at the correct Z level of the tissue of interest. Depending on the experimental goals, a small ROI for the camera and memory-based high frame-rate acquisition mode may be desired for capturing fast probe dynamics. In those cases, adjust exposure time and lighting angles/intensity to maximize contrast at high frame-rates. Note that lower resolution, wider field of view images should also be taken to assess the range of tissue deformation. Collect the data obtained in a measurement run and record the corresponding measurement program for downstream analysis.

### Consideration of measurement errors

The TiFM2.0 system offers significant improvements from TiFM1.0 in time synchronization and probe positioning accuracy. The main error consideration remains the spatial accuracy of *x*_T_, at the pixel with the sharpest contrast gradient with a resolution limit of ~0.5µm in ideal conditions, producing the largest error term in low signal-to-noise conditions such as thick tissues, combining errors along the segmentation and tracking pipeline. Due to general hardware limitations, voltage output is not a reliable measure of probe movement; the use of the interferometer (*x*_C_) marks a significant improvement in the tracking of piezo movement. Below we provide a list of noise sources and practices for minimizing these sources of technical variability.

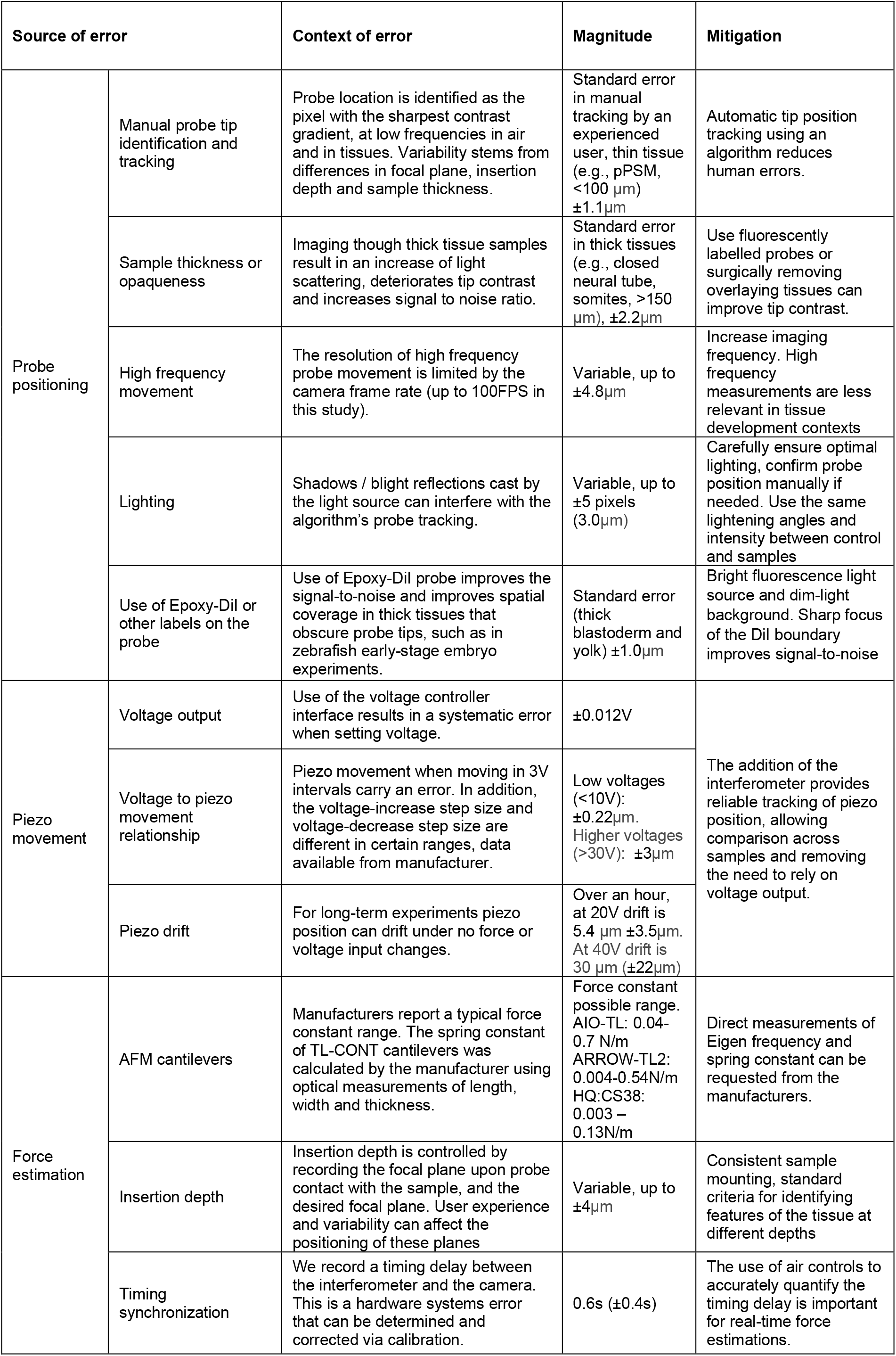

### Control samples

A semi-dilute polymer solution of 8.5%w/w, 13.5%w/w, 18.5% w/w and 23.5%w/w poly-isobutylene in hexadecane (PIB) (Sigma-Aldrich: 181455, H6703) was used. Hydrogel substrates were prepared following a protocol previously described^61,62^, where the stiffness of the hydrogel solutions was directly measured using Atomic Force Microscopy (AFM). Briefly, 150µL of the solutions described in the table below were sandwiched between two glass coverslips and left to polymerize for 20min. The gels were hydrated in PBS for 30 minutes, after which one coverslip was carefully removed, exposing the hydrogel. The hydrogel was then immobilized on a plastic 35mm dish using vacuum grease before TiFM measurements.

| Reagent | 79Pa | 369Pa | 1100Pa | 3130Pa | 5220Pa | Source |
|---|---|---|---|---|---|---|
| 40% Acrylamide [mL] | 0.75 | 0.75 | 0.75 | 1 | 1.875 | Bio-Rad 1610140 |
| 2% Bis-acrylamide [mL] | 0.18 | 0.3 | 0.5 | 1.125 | 0.375 | Bio-Rad 1610142 |
| PBS [mL] | 9.005 | 8.885 | 8.75 | 7.875 | 7.685 |  |
| To 1mL add: |  |  |  |  |  |  |
| TEMED [µL] | 15 | 15 | 1 | 1 | 15 | Thermofisher 17919 |
| 10% Ammonium persulfate [µL] | 50 | 50 | 10 | 10 | 50 | Fisher Scientific 50-995-775 |

### Rheometer measurements of control samples

The rheological measurements of PIB solutions were conducted on a rheometer (Anton Paar MCR302) using a cone plate (50 mm in diameter, 1° angle) at 25°C. The shear strain was set to 6% with a gap of 20μm, and the angular frequency was swept from 0.01 rad/s to 200 rad/s with 50 points per decade.

### Chicken embryos

Wild type chicken (*Gallus gallus*) eggs were supplied Medeggs Inc. Tg(CAG-GFP)^63^ and Tg(CAG-memGFP)^48^ chicken eggs were provided by the National Avian Research Facility (NARF) at University of Edinburgh. Eggs were stored in 15°C fridge and incubated in 37.5°C ~60% humidity incubators (Brisnea) for 24 hours for the HH4 early-stage experiment, ~40 hours to reach HH stage 10-12 for the body axis experiments, and 72 hours for the blood pressure experiment. Embryos were prepared using a modified EC culture^64^ protocol as described and incubated in a slide box lined with wet paper towels. Prior to TiFM experiments, embryos were further stabilized via a filter paper sandwich using enlarged windows to mitigate excess tension^40^ and mounted on a glass bottom image dish with a thin layer of albumen or agarose-albumen culture media^29^. A small amount of PBS or Ringer’s solution was added on top of the embryo prior to probe insertion for short-term experiments. For longer-term experiments, the water compensation pipette and pump were used at a flow rate of 8 µL/min, with a solution of 70% dH_2_O, 20% Ringers, 10% thin albumen. When the heating insert is used, the outer ring temperature was set at 42°C to allow the dish centre to reach ~36°C at the embryo location.

### Zebrafish embryos

Wild-type zebrafish AB and TL strains were used. Embryos were kept in 1X Embryo medium (E3 medium, reagents included: NaCl, KCl, CaCl_2_•H_2_O, MgSO_4_•7H_2_O, HEPES, ddH_2_O, pH to 7.4 with 10M NaOH) at 28.5°C. After dechorionation the embryos were kept in 1X E3 medium in a culture dish covered with agarose gel at 28.5°C. To hold the fish embryos and prevent rotations during measurements, a thin layer of 2% agarose was laid on the imaging dish, which was dug into with pipette tips to create two intersecting circles. E3 fish medium was then added to the imaging dish followed by loading the dechorionated embryo into one of the circles. The tissue to be tested is then rotated to face the opening between the 2 circles. The extra medium was then removed, leaving only a sufficient amount to cover the embryo. The probe was then inserted into the other circle, close to the test tissue. To ensure reliable tracking of the probe once it enters thicker tissue areas, a DiI-Epoxy bead was glued to the end of the cantilever to make a fluorescent trailing probe and imaged fluorescence. All procedures involving embryos were regulated by the Animals (Scientific Procedures) Act 1986, Amendment Regulations 2012, and in compliance with ethical standards at the University of Cambridge.

### Data analysis

Movies were analysed in Fiji (ImageJ^65^). To track the probe(s), the raw movies were cropped into small sections containing the range of the probe movement. Linear adjustments (Brightness and contrast) were applied to sharpen the probe vs sample contrast. These processed movies were then used as input together with the meta information file (time stamps) and the interferometer data output file in a custom analysis script, to obtain the tracks of the probe(s) and the holder positions for downstream analysis. To estimate forces on the probe, deflection was determined by comparing the tracks in the sample with tracks in the control (no-resistance) run, as described in the text. In some cases, manual tracking of probe location was performed from the kymograph generated in Fiji (ImageJ). For cell shape analysis, cell membranes were segmented using Cellpose^49^ and subsequently corrected manually.

## Supplemental Information

**Figure S1.**
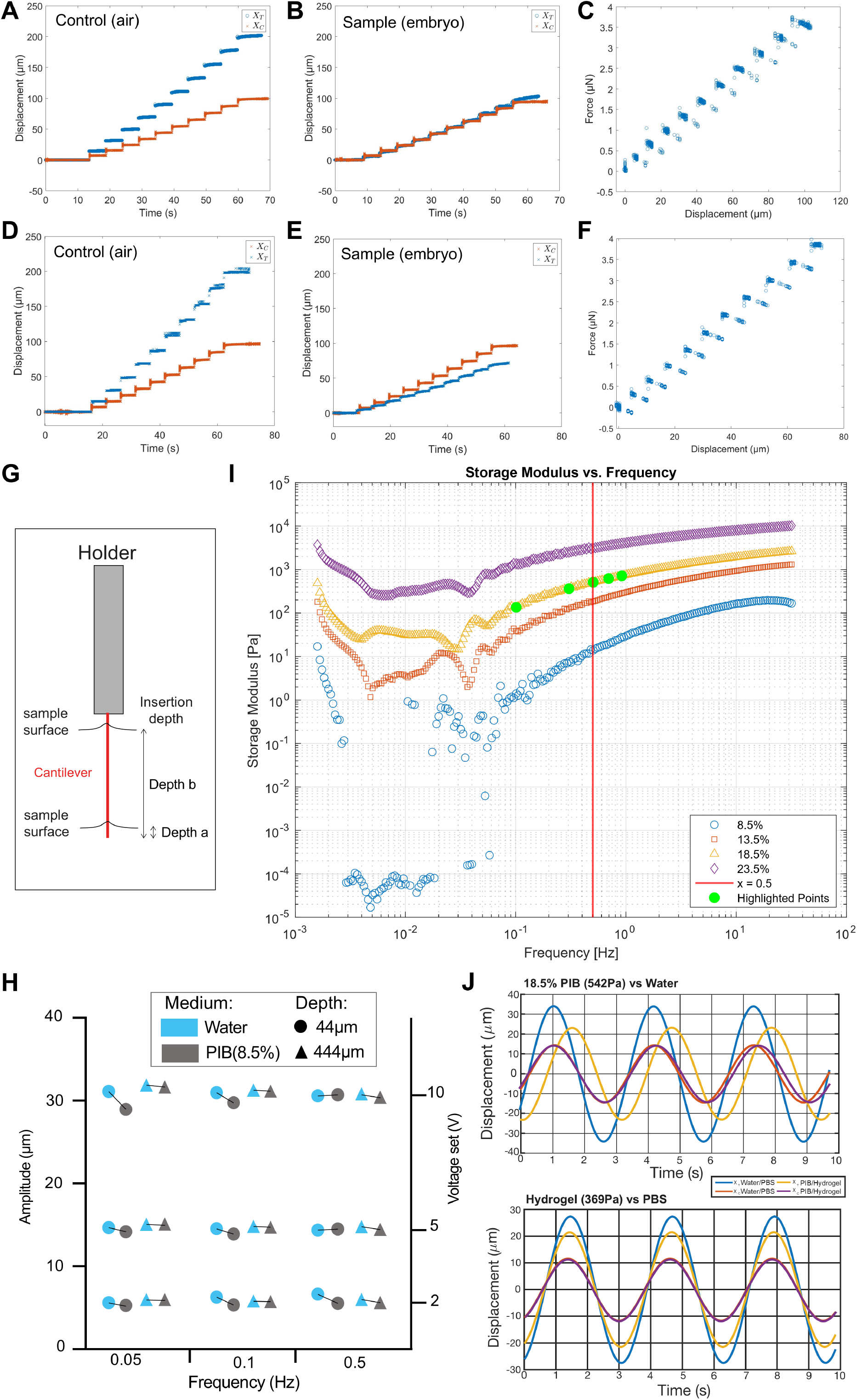
Additional supporting data. (A-F) Two additional repeats (A-C, D-F, respectively) related to Figures 1E-G. A,B,D,E show the original data of the interferometer positioning measurements (X_C_) and tracking of the probe tip (X_T_). C,F show the calculated force dynamics on the probe by comparing data in the control and embryo sample as described in the main text. (G) Diagram of single-probe mechanical tests at different depths and for different samples in Figure 7D and panel H. (H) Comparison of probe amplitude between water and PIB. Comparison between 3 different voltages (2V, 5V and 10V), over 3 different sinusoid frequencies (0.05Hz, 0.1Hz and 0.5Hz), at two different depths (circle: 44 µm and triangle: 444 µm). Amplitude calculated by fitting between 2 and 7 full periods of the sin wave. Probe force constant *k* = 0.31N/m. (I) Storage modulus (G’) [Pa] of 8.5%, 13.5%, 18.5% and 23.5% PIB, measured using a standard rheometer and plotted against the angular frequency of sweep. Line at x=0.5Hz represents reference values used in Figure 7B, green highlighted points show different frequencies used in Figure 7C. (J) Example comparison using the same driving program in water and 18.5% PIB (**i**, 10V max, 0.5Hz, *k* = 0.5N/m) and in PBS and 369Pa Hydrogel (**ii**, 10V max, 0.5Hz, *k* = 0.2 N/m). The curves plotted were fitted sine waves on the raw data traces, the interferometer traces of two samples were aligned so that the differences in amplitude and phase of the probe tip tracks are highlighted. The large phase lag between the interferometer signal (driver) and the probe tip in the PIB shows PIB’s high viscosity.

### Supplemental Movie Legends

**Movie S1**. Double probe (*k* = 0.2N/m) stretching in aPSM and pPSM. Visible insertion wounds can be tracked after probe retrieval. Anterior to the left. Timestamp in mm:ss. Probe retrieval at 5:02.

**Movie S2**. Cell shape changes under tension. HH4 mem-GFP+ anterior neural plate area was imaged through bidirectional stretching. Probe force constant *k* = 0.2N/m. Voltage controlling the double-motor was increased by 5V steps (to prevent tissue tearing) to eventually 40V and held for 1 hour. Timestamp shows minutes and seconds (imaging frequency was 2Hz). The movie shows the first 4 minutes of the experiment (a focus adjustment around 03:45). Probes can be seen exiting the field of view on the sides as the stepwise stretching continued. Tissue and cells can be seen to change shape in response to the imposed tension. Scale bar: 100µm.

**Movie S3**. Step-wise closing and opening of the neural tube using the double-probe setup. Probes are inserted into the ectoderm lateral to the neural folds. Anterior to the top. Timestamp in mm:ss.

**Movie S4**. Permanent deformation of axis curvature towards right side upon push and incubation (1hr) with notochord anchored probe. Compared to 1min push on the same embryo (HH11). Push to 40V in stepwise manner. Tracks seen in Figure 5H. Imaging frequency during push and after probe retrieval 1Hz (1 per second). Imaging frequency during incubation 0.0167Hz (1 per minute). Timestamp in hh:mm:ss.

**Movie S5**. Representative examples of raw imaging data of the probe tip in hydrogel samples. Probe force constant *k* = 0.2N/m, set to 10V and with a sinewave frequency of 0.5Hz, at an insertion depth of 230 µm. While the motor and probe holder move the same way in each sample, the probe movement can be seen to decrease as the stiffness increases. In very stiff samples, the probe is locally “stuck” and rotates around the half insertion point, resulting in the probe tip moving in an opposite direction to the motor (e.g., 3130Pa and 5220Pa). Timestamp in ss:ms.

**Movie S6**. Representative examples of raw imaging data of the probe tip in embryonic tissues. Probe force constant *k* = 0.2N/m, at 10V, sinewave frequency of 0.5Hz. Tissue insertion depth is approximately 70µm. While the motor and probe holder move the same way in each sample, the probe movement amplitudes are distinct. PSM, presomitic mesoderm. Timestamp in mm:ss:ms.

**Movie S7** Double probe oscillation in aPSM (anterior PSM) and pPSM (posterior PSM). Bright spot is the sending probe, dark line is the detecting probe. *k* = 0.03N/m, 10V, 0.5Hz. In addition to amplitude differences, differences in tissue deformation between probes can be observed. Timestamp in hh:mm:ss.

**Movie S8**. Labelled probe pushing different tissues in zebrafish embryos. The probe moves linearly towards a maximum distance driven by the motor (20V, 0.1 Hz). Fish medium (1st row), animal pole at 50% epiboly (2nd row) and dorsal trunk at ~15 somite stage (3rd row). Probe front is obscured by tissue after contact and cannot be further tracked. Tracking the trailing dye bead allows comparison of tissue resistance.

